# Mispacking of the F87 sidechain drives aggregation-promoting conformational fluctuations in the subunit interfaces of the transthyretin tetramer

**DOI:** 10.1101/2024.02.26.582172

**Authors:** Xun Sun, James A. Ferguson, Ke Yang, Robyn L. Stanfield, H. Jane Dyson, Peter E. Wright

**Author notes:** Equal contribution (Xun Sun, James A. Ferguson, Ke Yang).

## Abstract

Aberrant formation and deposition of human transthyretin (TTR) aggregates causes transthyretin amyloidosis. To initialize aggregation, transthyretin tetramers must first dissociate into monomers that partially unfold to promote entry into the aggregation pathway. The native TTR tetramer (T) is stabilized by docking of the F87 sidechain into an interfacial cavity enclosed by several hydrophobic residues including A120. We have previously shown that an alternative tetramer (T*) with mispacked F87 sidechains is more prone to dissociation and aggregation than the native T state. However, the molecular basis for the reduced stability in T* remains unclear. Here we report characterization of the A120L mutant, where steric hindrance is introduced into the F87 binding site. The X-ray structure of A120L shows that the F87 sidechain is displaced from its docking site across the subunit interface. In A120S, a naturally occurring pathogenic mutant that is less aggregation-prone than A120L, the F87 sidechain is correctly docked, as in the native TTR tetramer. Nevertheless, ^19^F-NMR aggregation assays show an elevated population of a monomeric aggregation intermediate in A120S relative to a control containing the native A120, due to accelerated tetramer dissociation and slowed monomer tetramerization. The mispacking of the F87 sidechain is associated with enhanced exchange dynamics for interfacial residues. At 298 K, the T* populations of various naturally occurring mutants fall between 4–7% (Δ*G* ∼ 1.5– 1.9 kcal/mol), consistent with the free energy change expected for undocking and solvent exposure of one of the four F87 sidechains in the tetramer (Δ*G* ∼ 1.6 kcal/mol). Our data provide a molecular-level picture of the likely universal F87 sidechain mispacking in tetrameric TTR that promotes interfacial conformational dynamics and increases aggregation propensity.

## Introduction

Transthyretin (TTR) is an abundant tetrameric protein that functions as a transporter for thyroxine in blood. Aggregation of TTR leads to TTR amyloid disease, including wild-type transthyretin cardiac amyloidosis and familial cardiomyopathy and polyneuropathy associated with hereditary mutations (1). The TTR aggregation pathway begins with dissociation of the TTR tetramer to form monomers (2) or unstable dimers that rapidly dissociate (3). Wild-type (WT) transthyretin cardiac amyloidosis affects up to 25% of the population with age over 80. Pathogenic mutations that weaken the TTR tetramer are associated with earlier ages of onset of TTR amyloidosis. Thus, it is important to understand how the TTR tetramer becomes destabilized to facilitate dissociation.

We have previously shown using ^19^F-NMR that an alternatively-packed TTR tetramer (T*) state is more prone to dissociation and aggregation than the native TTR tetramer (designated the T state) (4). In the native TTR tetramer, the F87 sidechain packs into a hydrophobic pocket in the neighboring subunit across the strong dimer interface (Figure 1A), contributing to the stability of the tetramer. The pocket is lined by several hydrophobic residues, including A120 (Figure 1B). By contrast, the F87 sidechain is mispacked in the T* state, which in native human TTR has a population of ∼7%. The low population of the T* state makes structural characterization challenging. To circumvent this problem, we have shown that the T* population can be increased by mixing WT-TTR with a designed A120L mutant; steric clash arising from the bulky Leu sidechain forces the F87 sidechain to adopt an alternative conformation. In this work, we use A120L as a model for the T* state and solve its X-ray structure to highlight how the F87 sidechain is mispacked in the T* state. We also study a naturally occurring A120S mutant that is more stable than A120L but less stable than the wild-type TTR. Using both ^19^F and amide NMR probes, we show that the conformational dynamics are enhanced in several interfacial residues in the T* state, which is likely linked with its higher aggregation propensity.

**Figure 1.**
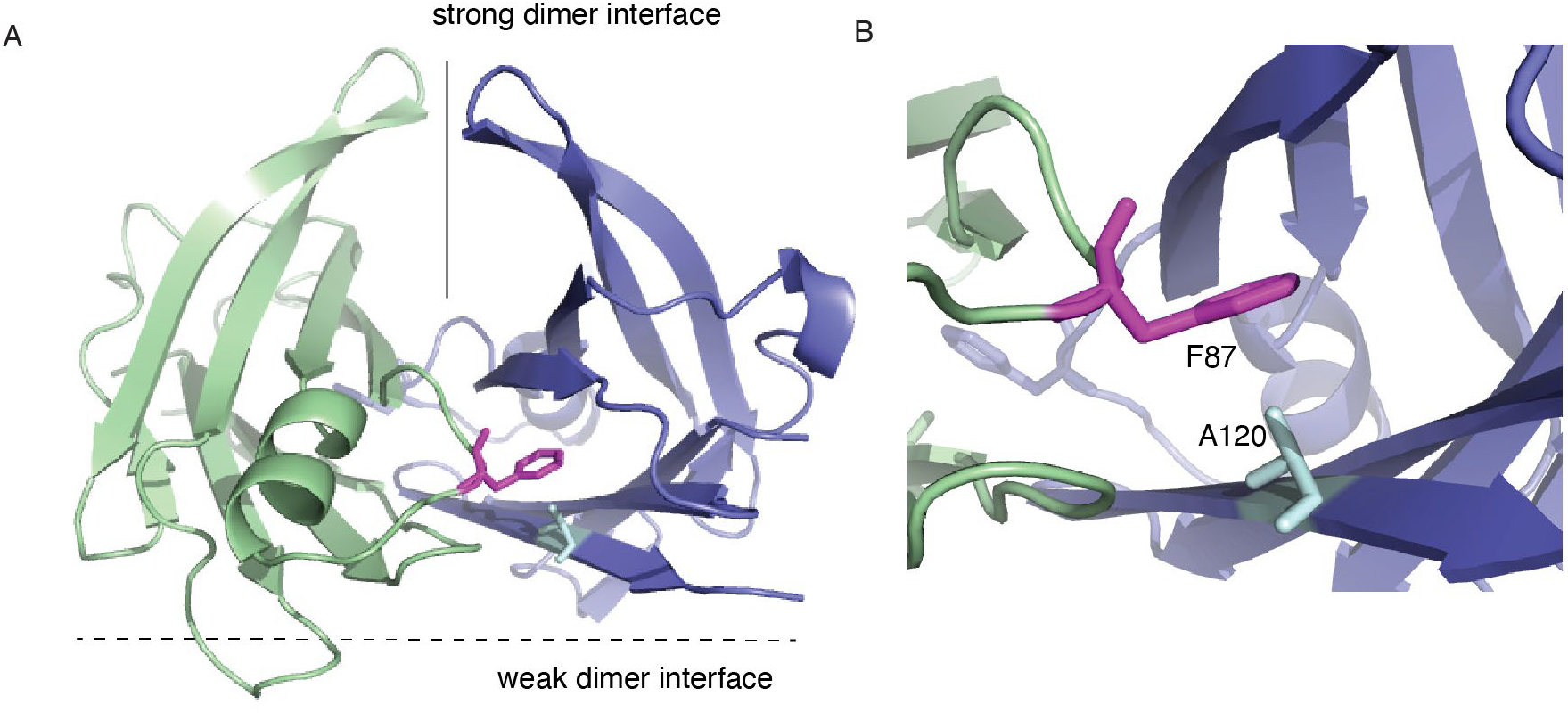
The F87 sidechain (magenta) packs into a hydrophobic pocket in the neighboring subunit across the strong dimer interface in a WT TTR structure (PDB: 5CN3) shown in a global view (A) and a closeup view (B).

## Materials and Methods

### Protein expression and purification

WT human TTR and the A120S and A120L TTR variants were expressed and purified as previously reported (5). Purification of Cys-derived oligomers based on C10S-S85C or C10S-S100C and labeling for ^19^F NMR with 3-bromo-1,1,1-trifluoroacetone (BTFA) was performed as previously described (6). Unless noted, all experiments were performed in 10 mM potassium phosphate, and 100 mM KCl at pH 7.0 (NMR buffer). Uniformly ^15^N labeled WT TTR was expressed in M9 medium containing 1 g/L (^15^NH_4_)_2_SO_4_ and 3 g/L ^12^C-glucose.

### Size exclusion chromatography

A Superdex 75 gel filtration column was pre-equilibrated in NMR buffer. A120L or wild-type TTR was then loaded at a flow rate of 0.7 mL/min.

### ^19^F-NMR spectroscopy

^19^F NMR spectra were recorded at 298 K using Bruker Avance 600 or Bruker Avance 700 MHz spectrometers as described previously (6). The fitting of *K*_d_ for the T ⇌ 4M equilibrium for A120S was performed as previously described (7).

### Real-time NMR mixing experiments

^15^N labeled WT TTR and the unlabeled A120L variant were mixed at a 3:1 molar ratio (WT:A120L = 600 μM: 200 μM) or a 1:1 molar ratio (WT:A120L = 200 μM: 200 μM) in NMR buffer with 10% D_2_O added for the lock signal. At these concentrations, A120L is a mixture of tetramer and monomer (4). Standard [^1^H,^15^N]-heteronuclear single quantum coherence spectra (HSQC) were acquired using Bruker Avance 600 or 700 MHz spectrometers as previously described (8). In the 3:1 mixing experiment, for each delay point in the indirect ^15^N dimension, 12 scans were collected, giving a total acquisition time of 60 min per 2D spectrum. A total of 72 2D spectra were recorded over 3 days. In the 1:1 mixing experiment, 32 scans were collected for each ^15^N delay and a total of 45 spectra were recorded for 5 days. The backbone amide assignments were transferred from those of wild-type human TTR (BioMagResBank code 27514) (9). The peak intensities of all non-overlapped peaks were extracted by using CcpNmr (10) and normalized to that of the C-terminus E127. For peaks with pronounced intensity decays, a global fit to a single-exponential function was performed using the MATLAB (R2022b) function *nlinfit* to extract plateau intensities. The error in the fitted rate constant was estimated as one standard deviation with 68% confidence level using *nlpredci*. All NMR data processing and analysis were carried out on the NMRBox platform (11).

### A120L and A120S X-ray crystallography

Crystallization screens were set up using the sitting drop vapor diffusion method with the A120L and A120S mutants at a concentration of 9–10 mg/mL in NMR buffer. The crystallization condition for the A120L trial was 0.1 M sodium cacodylate (pH 6.5), 1M sodium chloride, 10% (v/v) glycerol, and 30% (v/v) PEG 600 at 293 K. The cryoprotectant was 30% (v/v) PEG. The crystallization condition for the A120S trial was 0.1 M MES at pH 6.0, 10% (v/v) glycerol, 5% (w/v) PEG 1000, and 30% (v/v) PEG 600 at 293 K. The data were processed using HKL-2000 (12). Molecular replacement was performed using Phaser (13) with the TTR structure (PDB ID: 2ROX) as a search model. Models were refined using phenix.refine (14), refmac (15), and Coot (16). The coordinates have been deposited in the Protein Data Bank with accession codes 8W2W (A120L) and 8W1N (A120S).

### Molecular dynamics (MD) simulations

The X-ray structures of wild-type human TTR (PDB: 5CN3) and A120L (PDB: 8W2W) were used as a starting model for canonical ensemble MD simulations with explicit solvent using the AMBER ff14SB force field (17) and AMBER 16 software (18) as previously described (3). The X-ray crystallographic symmetry was used to create a tetrameric TTR model as an initial conformation using PyMOL (2.5.0) as described previously (5) and one 400-ns MD simulation for each tetramer was run using protocols as previously described (3). The mass-weighted root-mean-square-fluctuations (RMSF) per residue (19) for backbone heavy atoms were extracted using CPPTRAJ (20), averaged over all four chains of the tetramer, and analyzed using MATLAB. The surface area of the F87 sidechain was estimated using VADAR (21) based on the wild-type structure (PDB: 5CN3).

## Results

### A120S is more stable than A120L but less stable than wild-type TTR

To measure the relative populations of monomer (M) and tetramer (T), we constructed variants of the C10S-S85C-BTFA (TTR^F^) construct, which has distinct ^19^F chemical shifts for the monomeric and tetrameric species (6). We introduced both the A120S and A120L mutations into TTR^F^ separately and denote the resulting constructs as A120S^F^ and A120L^F^, respectively. Figure 2A shows that A120S^F^ forms a mixture of 88% tetramer (T) and 12% monomer (M) at 10 μM concentration. Fitting the populations of T and M from a series of A120S^F^ spectra over a range of concentrations (Figure 2B), we determined the *K*_d_ for the T ⇌ 4M equilibrium to be 6.5 x 10^-19^ M^3^, comparable to the *K*_d_ = 6.5 x 10^-19^ M^3^ for K80D^F^ (7) and much weaker than the *K*_d_ = 9 x 10^-25^ M^3^ of wild-type human TTR (22). By contrast, A120L^F^ is fully monomeric at 8 μM as shown by ^19^F-NMR (Figure 2A), consistent with gel-filtration analysis (Figure S1). These results show qualitatively that A120L is thermodynamically less stable than A120S; however, we were unable to determine a tetramer/monomer *K*_d_ for A120L^F^ because it aggregates at the higher concentrations needed for tetramer formation.

**Figure 2.**
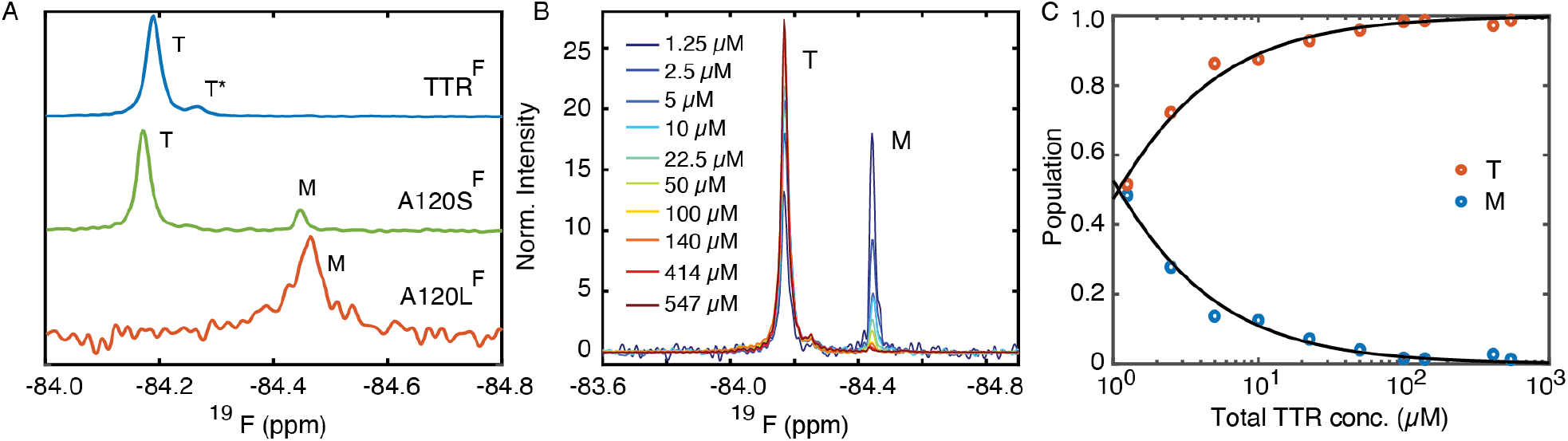
(A) ^19^F-NMR spectra of TTR^F^-based mutants at 298 K and pH 7.0 showing the ^19^F chemical shifts of the native tetramer (T), the alternative tetramer (T*) and the monomer (M). The concentration of A120L^F^ was 8 μM and the other constructs were at 10 μM. (B) ^19^F-NMR concentration titration for A120S^F^ at 298 K and pH 7.0. Spectra were normalized by total peak areas. (C) The fitted apparent *K*_d_ = 6.5 x 10^-19^ M^3^ for the T ⇌ 4M equilibrium was determined by fitting the relative population of T and M from (B).

As indicated by the tetramer/monomer *K*_d_, A120S is thermodynamically less stable than wild-type TTR. We next measured the acid-mediated aggregation kinetics of A120S^F^ at pH 4.4 (Figure 3A–B) and analyzed the data using a three-state T ⇌ M ⇌ Aggregates (A) model (6). The tetramer dissociation rate constants are greater for A120S^F^ than in the control TTR^F^ at 298 and 310 K (Table 1). Notably, the reverse tetramerization rate constant of A120S^F^ is 7-fold slower than TTR^F^ at 298 K. Accordingly, we observed a higher level of the monomeric aggregation intermediate (M) in A120S^F^ than in TTR^F^ (Figure 3C). We have previously shown (6) that mutation of the F87 sidechain to Ala also enhances the kinetics of tetramer dissociation, slows reassembly, and increases the steady population of the aggregation intermediate M (Figure 2C, Table 1). Taken together, our data indicate that disruption of the hydrophobic interaction between the sidechains of F87 and A120, by mutation of either residue, destabilizes the tetramer and increases the aggregation propensity (Table 1).

**Table 1.** ^19^F-NMR aggregation kinetics based on the T ⇌ M ⇌ A model. The results for TTR^F^ and F87A^F^ were taken from Ref. (6) for comparison with A120S^F^. The slow relaxation rate constant γ_2_ describes the aggregation propensity.

| Mutant | Temp. (K) | $k_1$ (h<sup>-1</sup>) | $k_{-1}$ (h<sup>-1</sup>) | $k_2$ (h<sup>-1</sup>) | $k_{-2}$ (h<sup>-1</sup>) | $\gamma_2$ (h<sup>-1</sup>) |
| --- | --- | --- | --- | --- | --- | --- |
| TTR<sup>F</sup> | 310 | 0.10 ± 0.01 | 0.75 ± 0.12 | 0.73 ± 0.03 | 0.03 ± 0.01 | 0.06 ± 0.01 |
| TTR<sup>F</sup> | 298 | 0.13 ± 0.02 | 0.63 ± 0.10 | 0.06 ± 0.01 | 0.01 ± 0.01 | 0.02 ± 0.01 |
| A120S<sup>F</sup> | 310 | 0.22 ± 0.03 | 0.54 ± 0.14 | 0.93 ± 0.09 | 0.07 ± 0.01 | 0.16 ± 0.01 |
| A120S<sup>F</sup> | 298 | 0.24 ± 0.04 | 0.06 ± 0.02 | 0.05 ± 0.01 | 0.02 ± 0.01 | 0.05 ± 0.01 |
| F87A<sup>F</sup> | 310 | 1.60 ± 0.16 | 0.18 ± 0.04 | 0.56 ± 0.01 | 0.09 ± 0.01 | 0.57 ± 0.02 |
| F87A<sup>F</sup> | 298 | 0.46 ± 0.02 | 0.04 ± 0.01 | 0.06 ± 0.01 | 0.03 ± 0.01 | 0.08 ± 0.01 |

**Figure 3.**
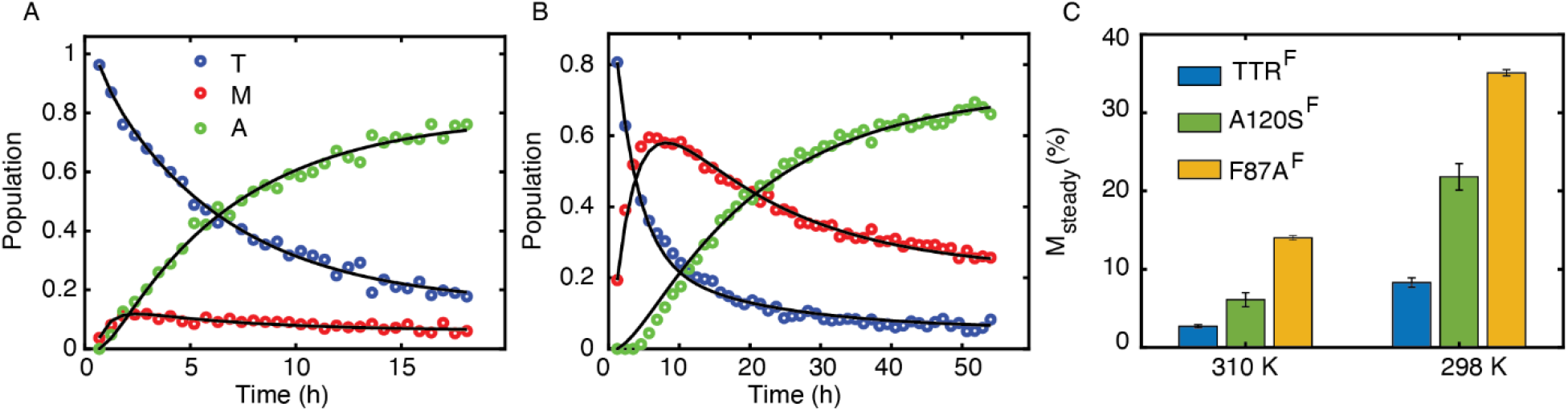
(A–B) ^19^F-NMR aggregation assays at pH 4.4 for A120S^F^ at 310 K (A) and 298 K (B). The black lines denote the fits from using a three-state aggregation kinetics model T ⇌ M ⇌ A where T, M, and A stand for tetramer, monomeric intermediate, and aggregates, respectively. (C) The steady-state monomer concentration (M) of TTR^F^, A120S^F^, and F87A^F^ at 310 and 298 K. The data for TTR^F^ and F87A^F^ were replotted from Ref. (6).

### The F87 sidechain is mispacked in the X-ray structure of A120L but not A120S

To understand the molecular basis of the distinct F87-A120 interactions in A120S and A120L, we next solved their X-ray structures. The A120L and A120S constructs crystallized in space groups I 2 2 2 and P2_1_2_1_2, with one and two TTR protomers in the asymmetric unit, respectively (Figure 4A). The diffraction data were refined to 2.07 and 1.60 Å, respectively (Table S1). The mean Cα RMSD values for both X-ray structures relative to a wild-type TTR structure (PDB: 5CN3) are less than 0.5 Å. This comparison shows that the mutations do not significantly perturb the tertiary structure of TTR tetramers (Figure S2A–B). The B-factor profile of A120S closely resembles that of the wild-type human TTR whereas A120L displays elevated B factors throughout the EF region (Figure 4B).

**Figure 4.**
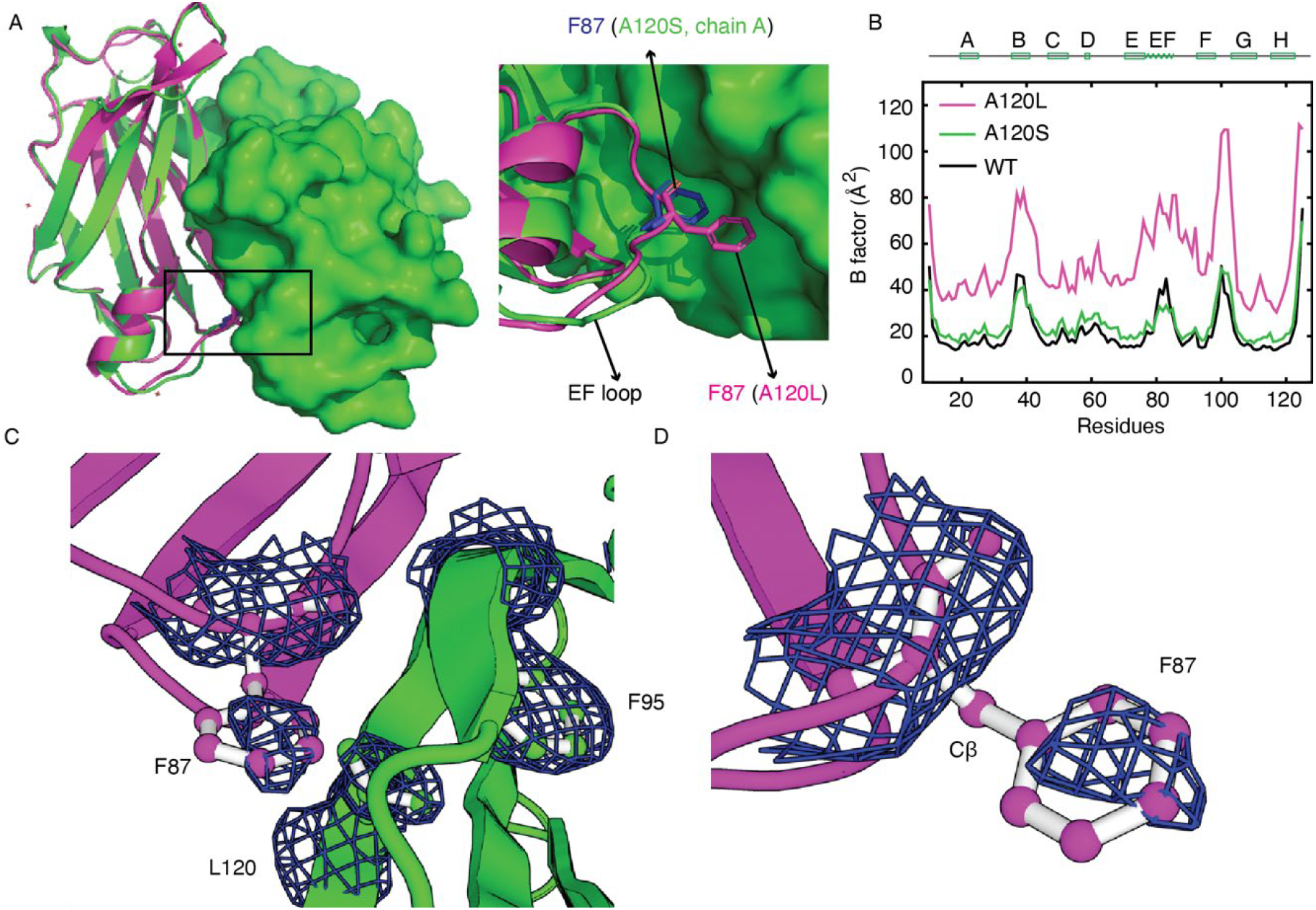
(A) X-ray crystal structures of A120S chain A (green) and A120L (magenta) shown as cartoons, with chain B of A120S shown as a surface model in a strong dimer. Inset: Close-up view of the structure, zoomed from the black box in (A), showing the distinct orientations of the F87 sidechain in A120L and A120S. (B) Comparison of mass-weighted crystallographic B factors for backbone heavy atoms averaged for all subunits in one unit cell for A120L and A120S X-ray structures, in comparison with the wild-type TTR (black, PDB: 5CN3). B factors were mass-weighted per residue and averaged over all four chains of the tetramer. (C) Sidechains of F87, F95 and L120 modeled into the electron density in the A120L dimer structure imposed by crystallographic symmetry. (D) F87 with the electron density map for the A120L monomer structure. The displayed maps are |2F_o_-F_c_| with σ = 1.00.

The difference in the orientation of the F87 sidechain between the wild-type and A120S TTR structures is subtle (Figure S2C) while the F87 sidechain is displaced from its binding pocket and occupies a solvent-exposed position in the A120L structure (Figure 4C, S2D). In the A120L structure, the sidechain of L120 occupies the hydrophobic pocket that accommodates the F87 sidechain in the wild-type and A120S structures (Figure S2C–D). The distinct orientations of the F87 sidechains in A120L and A120S are consistent with the greater volume of the Leu sidechain compared to Ser; occupancy of the binding pocket by the bulky Leu sidechain precludes docking of the F87 aromatic ring and explains the lower thermodynamic stability of A120L^F^ compared to A120S^F^ (Figure 2A). The electron density for the F87 sidechain in the A120L structure does not completely encompass the phenyl ring (Figure 4D). In addition, there is no electron density for the Cβ atom of F87, suggesting that multiple conformations are sampled by the F87 sidechain of the A120L mutant, a model of the T* state. Consistent with this, enhanced nanosecond timescale fluctuations of the F87 sidechain about the χ1 and χ2 dihedral angles were observed in all-atom MD simulations of the A120L tetramer, but not in simulations of wild-type TTR (Figure 5A–D).

**Figure 5.**
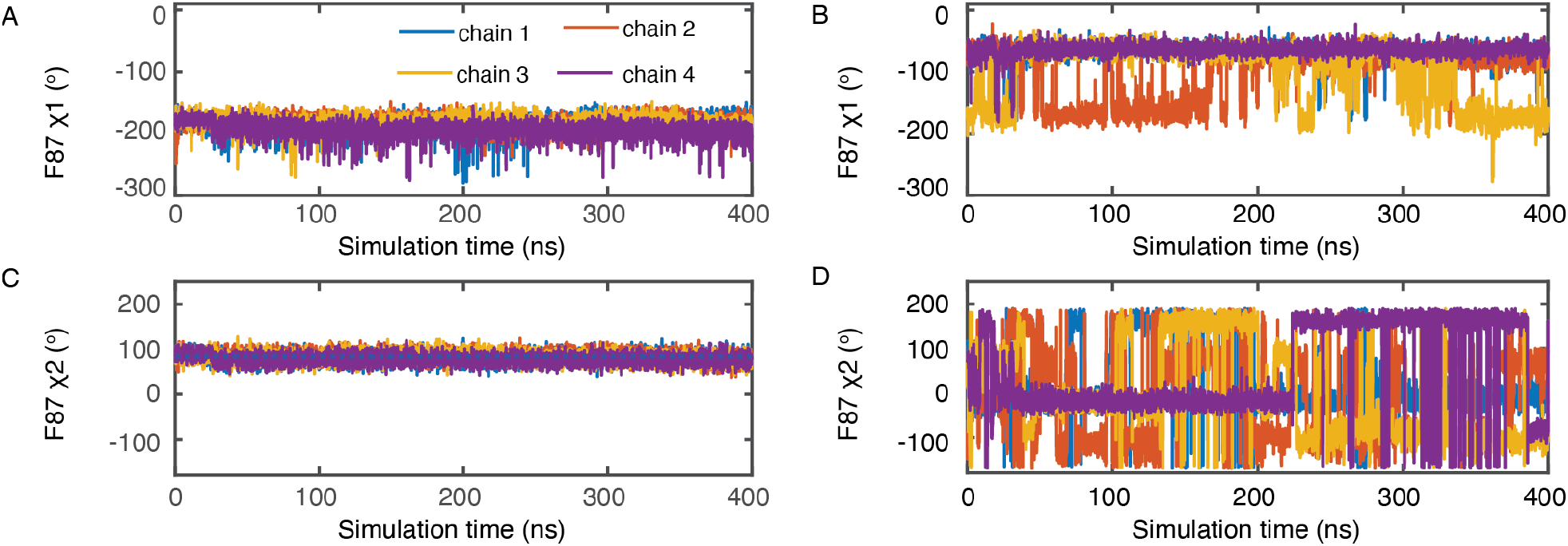
Nanosecond dynamics of the F87 χ1 (A–B) and χ2 (C–D) dihedral angles in canonical ensemble MD simulations for WT (A, C) and A120L tetramers (B, D).

### The T* population is conserved and consistent with displacement of a single F87 sidechain within the tetramer

We used ^19^F-NMR to measure the population of the T* state in the pathogenic variants A25T, V30M, L55P, A81T, G83R, A97G, T119M, A120S, V122I (Table 2). All mutations were introduced into the TTR^F^ construct and coupled to BTFA as described previously (6). The T* population was determined from the relative intensities of the T and T* peaks. Including the T* population (7%) previously reported for TTR^F^ (4), all T* populations are within a range of 4–7%, corresponding to Δ*G* relative to the correctly packed tetramer T of ∼ 1.5–1.9 kcal/mol at 298 K, based on the Boltzmann distribution. The buried surface area of F87 decreases from 120 Å^2^ in the WT tetramer to 11 Å^2^ in A120L, where the aromatic ring is displaced from its binding pocket and becomes exposed to solvent. Based on a coefficient of 15 cal/mol/ Å^2^ at 298 K (23), the associated Δ*G* is ∼ 1.6 kcal/mol. This comparison indicates that the measured T* population is a result of the Boltzmann distribution associated with the energetic difference linked to undocking of one of the four F87 sidechains from its binding pocket in the strong dimer interface of the tetramer.

**Table 2.** Population of the T* state for TTR variants at 298 K and pH 7.0. The superscript ^F^ denotes that the mutations were introduced into the C10S-S85C-BTFA (TTR^F^) background. The T* population for TTR^F^ from Ref. (4) is included for comparison.

| Mutant | T* (%) |
| --- | --- |
| TTR<sup>F</sup> | 6.8 ± 1.3 |
| A25T<sup>F</sup> | 5.1 ± 0.7 |
| V30M<sup>F</sup> | 6.4 ± 0.7 |
| L55P<sup>F</sup> | 6.7 ± 0.2 |
| A81T<sup>F</sup> | 7.4 ± 0.5 |
| G83R<sup>F</sup> | 5.4 ± 2.1 |
| A97G<sup>F</sup> | 4.4 ± 2.3 |
| T119M<sup>F</sup> | 4.7 ± 0.7 |
| A120S<sup>F</sup> | 3.5 ± 2.2 |
| V122I<sup>F</sup> | 4.7 ± 0.6 |
| C10S-S100C<sup>F</sup> | 4.1 ± 0.3 |
| Mean | 5.4 ± 1.3 |

### The T* state displays enhanced conformational fluctuations of interfacial residues

We have so far relied on one ^19^F probe, the CF_3_ group of BTFA coupled to C10S-S85C TTR, to report on the T* state. To expand the number of probes using backbone amide cross peaks, we mixed ^15^N-labeled wild-type TTR with unlabeled A120L and recorded a series of HSQC spectra over 5 days at 298 K (Figure 6A). We observed extensive intensity decay for a set of amide cross peaks, which could be fitted by a global rate constant (0.036 ± 0.003 h^-1^) (Figure 6B, S3). The population of the T* state, as represented by the amplitude loss of amide cross peaks, increases as the molar ratio of A120L:WT is changed from 1:3 to 1:1 (Figure 6C). Interfacial residues, particularly at the AB loop, and F and H β-strands, show pronounced peak broadening together with residues in the EF helix and across the entire DAGH β-sheet (Figure 7A–B).

**Figure 6.**
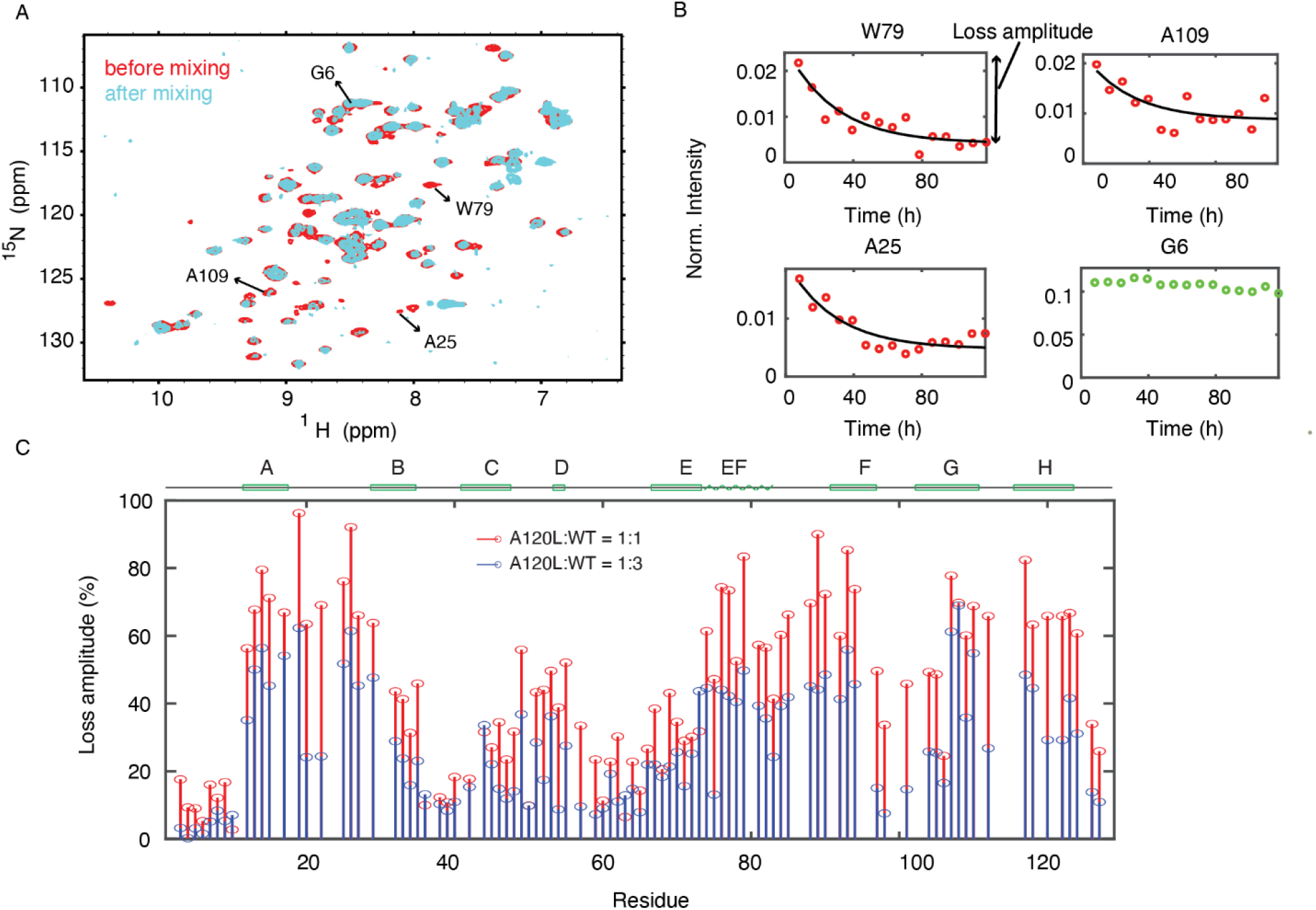
(A) ^15^N HSQC spectra of ^15^N-labeled WT TTR before and immediately after mixing with unlabeled A120L at a 1:1 molar ratio. (B) Time-dependent peak intensity changes of the three residues that show the largest loss of signal amplitude, normalized by the peak intensity of E127 at the first time point. The black solid lines denote single exponential fits. For comparison, the G6 amide shows nearly constant peak intensity (green data points). The cross peaks for these residues are identified by arrows in the HSQC spectrum in panel (A). (C) The percentage of cross peak amplitude loss after 5 days as a function of amino acid residue number at 1:1 (red) and 1:3 (blue) A120L:TTR molar ratios. The location of the secondary structure elements is indicated at the top of (C).

**Figure 7.**
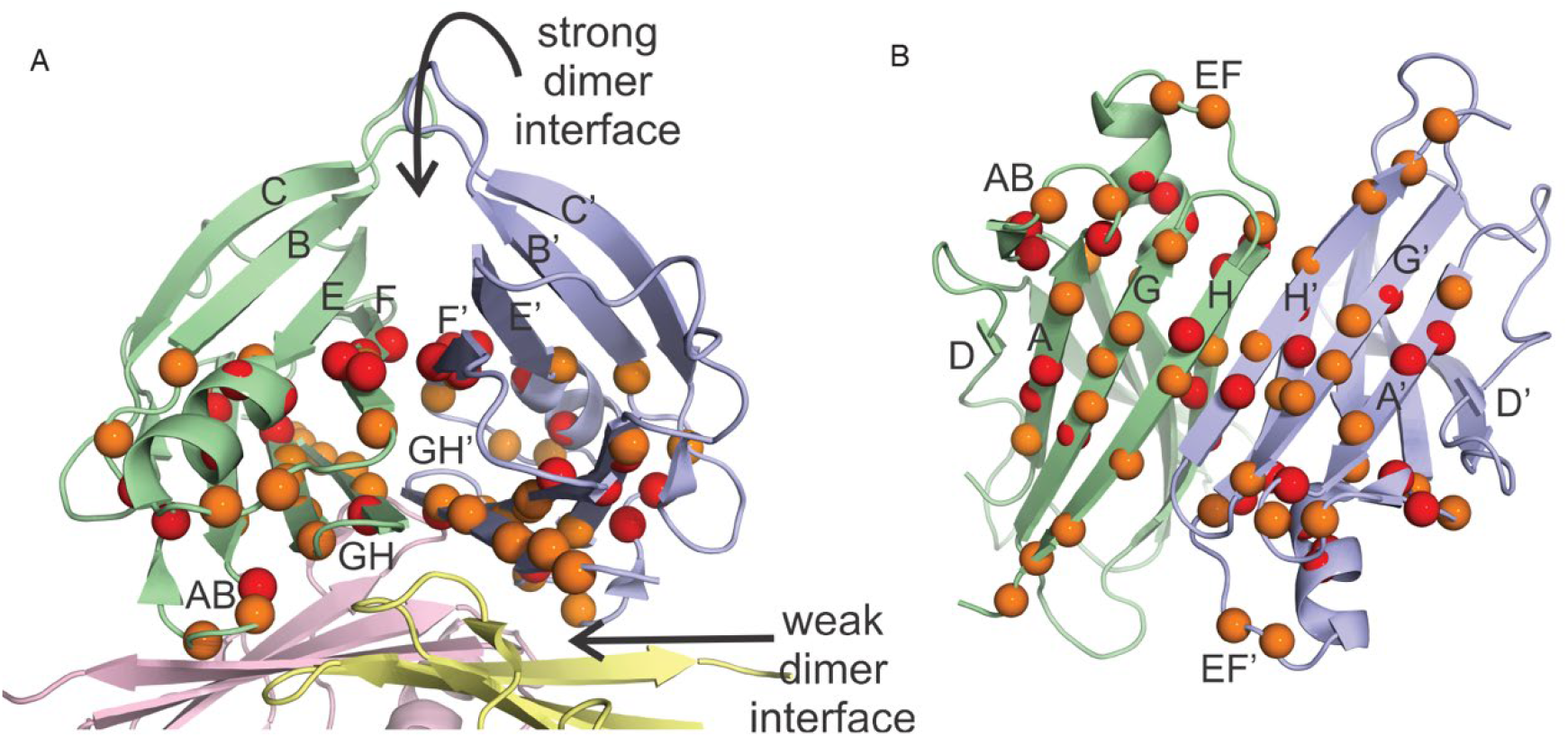
(A) Location of residues that exhibit greater than 70% (red spheres) or 60-70% (orange spheres) HSQC cross peak intensity loss upon mixing unlabeled A120L and ^15^N-labeled WT TTR in a 1:1 ratio. The structure shown is that of a WT tetramer (PDB code 5CN3). (B) The DAGH β-sheets in the strong dimer as viewed from the weak dimer interface. The residues that exhibit cross peak intensity loss in the HSQC spectrum of the 1:1 mixture are colored as in panel (A).

## Discussion

The low population of the alternatively packed T* state (∼7%) in WT human TTR make the determination of its molecular structure challenging. We have previously shown that the population of the T* state of TTR^F^ can be increased by mixing with A120L and that the less stable T* state is associated with the mispacking of the F87 sidechain (4). In the present work, we determined the X-ray structure of the A120L tetramer as a model of the T* state. Due to steric clash with the bulky L120 sidechain, the F87 aromatic ring is displaced from its hydrophobic pocket in the neighboring subunit across the strong dimer interface of the tetramer and rotates into a solvent-accessible position (Figure 4C). By contrast, the F87 sidechain remains docked in the A120S structure (Figure S2C), explaining the enhanced thermodynamic stability of A120S compared to A120L at pH 7.0 (Figure 2A). The B factors of A120S are very similar to those of WT TTR whereas F87 displays elevated B factors throughout the EF region that likely reflect enhanced dynamics due to loss of the F87 sidechain packing interactions. We also note that residues in the FG loop (residues 97–104) consistently show elevated B factors in both the A120L and A120S structures as well as in wild-type TTR (Figure 4B).

Displacement of F87 from its binding pocket, together with increased backbone flexibility in the EF region, allows the sidechain to sample multiple solvent-exposed conformations. In the MD simulations for A120L, F87 samples multiple χ1 and χ2 dihedral angles (Figure 5B, D), suggesting that in the T* state, F87 can interconvert between these conformations on the nanoscale timescale. By contrast, the direct conversion between the native T and the alternative T* state is much slower for TTR^F^ (0.038 h^-1^) (4) and is comparable to the subunit exchange rate constant of the wild-type TTR (0.02-0.04 h^-1^) (24; 25). Consistently, the rate at which the amide cross peaks lose intensity when wild-type TTR is mixed with A120L (0.036 h^-1^) is also within this range of subunit exchange rates (Figure 6B, S3). The similarity in rate constants indicates that subunit exchange mediates direct exchange between the T and T* states of the wild-type TTR tetramer. Moreover, the loss of amide cross peak intensity upon mixing of ^15^N-labeled WT TTR with unlabeled A120L shows that the T* state is associated with increased conformational dynamics. Not all residues lose cross peak intensity to the same extent and interfacial residues, in particular, lose much of their cross peak intensity during the 5 days of data acquisition, indicating enhanced conformational fluctuations associated with the destabilized tetramer interfaces (Figure 6C, 7A, B). The amide cross peak intensity loss increases as the molar ratio of A120L mixed with wild-type TTR increases (Figure 6C), in agreement with the titration assay for TTR^F^ with A120L by ^19^F-NMR (4).

The interfacial residues that display enhanced conformational fluctuations and cross peak broadening in the T* state (Figure 7A–B) have broadened resonances in the unstable and pathogenic A25T variant (3). These interfacial residues are also perturbed due to the incorporation of 6-fluoro-tryptophan at W79 in the EF helix, which reduces tetramer stability (5), and many of undergo millisecond timescale conformational fluctuations revealed by NMR relaxation dispersion studies of wild-type TTR and the V30M, L55P, and V122I variants (9). This comparison suggests that increased dynamics associated with perturbation of the protomer interfaces are coupled to the enhanced aggregation propensity of the T* state. The relevant dynamics are likely on the millisecond timescale, not on the fast nanosecond timescale probed in the MD simulations. Indeed, most of the residues that exhibit cross peak broadening in the T* state undergo millisecond time scale conformational fluctuations in monomeric TTR variants (26; 27), highlighting the importance of correctly packed subunit interfaces for stabilization of the TTR tetramer.

The observation that the population of the T* state is relatively conserved (∼4-7%) in WT human TTR and several pathogenic variants (Table 2) provides new insights into tetramer assembly. We have shown here and elsewhere (4) that TTR assembles to form both a correctly packed (T) and a mispacked tetramer (T*) in which an F87 aromatic ring is displaced from its binding pocket in the neighboring subunit across the strong dimer interface. The population of T* is consistent with a Boltzmann distribution over two tetrameric states (T and T*) that differ in free energy by ∼1.6 kcal.mol^-1^ at 298 K, the free energy associated with displacement and solvent exposure of a single F87 sidechain. This free energy change is identical to the ΔΔG between the T and T* states at 298 K for a destabilized A25T mutant determined by van’t Hoff analysis (3). Compared to a free energy change of 32.8 kcal.mol^-1^ for WT TTR to fully dissociate from tetramer to monomers at 298 K (22), the free energy difference between the T and T* states is relatively small. We envisage that, during the process of tetramer formation, a small population of molecules assemble with a misplaced F87 sidechain, i.e. form the T* state. Interconversion between the T and T* states is slow (0.038 h^-1^in TTR^F^) (4), so, once formed, the mispacked T* tetramer is kinetically trapped. Over long time periods, TTR will reach a thermodynamic equilibrium in which there is a Boltzmann distribution over the T and T* states, and likely even higher energy states in which more than one F87 phenyl ring is displaced from the strong dimer interface. However, the population of such postulated higher energy states is too small to be observable in our experiments.

In conclusion, we have used a combination of solution NMR, X-ray crystallography, and MD simulations to elucidate the structural consequences of F87 sidechain mispacking in the T* state, which is less stable than the native T state. Population of the T* tetramer is a natural consequence of the small energy difference between the native tetramer and one in which a single F87 aromatic ring is undocked from the strong dimer interface, resulting in a Boltzmann distribution of the T and T* states. The enhanced dynamics at interfacial residues likely predispose the T* TTR tetramers to dissociate more readily and subsequently enter the aggregation pathway.

## Data availability

The X-ray structures of A120L and A120S have been deposited in the PDB with accession codes as 8W2W and 8W1N, respectively.

## Acknowledgements

We thank Ben Leach for providing the untagged A120L plasmid, Maria Martinez-Yamout for expert advice in molecular biology, Gerard Kroon for expert assistance in NMR experiments, and Euvel Manlapaz for technical support.

