## Supplementary Material for "Mispacking of the F87 sidechain drives aggregation-promoting conformational fluctuations in the subunit interfaces of the transthyretin tetramer"

| Mutant | A120S (8W1N) | A120L (8W2W) |
|---|---|---|
| **Data Collection** | | |
| **Beamline** | ALS 5.0.3 | APS 23IDB |
| **Wavelength (Å)** | 0.9765 | 1.0332 |
| **Resolution (Å)** | 38.6-1.60 (1.63-1.60) | 32.0-2.07 (2.14 – 2.07) |
| **Space Group** | $P\ 2_1\ 2_1\ 2$ | I 2 2 2 |
| **Unit Cell (Å)** | a=43.11, b=86.28, c=64.09 | a=42.14, b=64.07, c=84.63 |
| **Total reflections**[a] | 240,058 (11,432) | 74382(675) |
| **Unique reflections** | 31,452 (1493) | 7206 (361) |
| **Redundancy** | 7.6 (7.6) | 10.3 (8.5) |
| **Completeness (%)** | 97.7 (96.1) | 99.7 (98.6) |
| **<I>/<σI>** | 20.7 (1.0) | 25.8 (3.2) |
| **$R_{merge}$ (%)**[b] | 9.4 (223) | 6.8 (62.8) |
| **$R_{meas}$ (%)**[c] | 10.2 (240) | 7.5 (70.3) |
| **$R_{pim}$ (%)**[d] | 3.7 (85.5) | 3.1 (31.0) |
| **$CC_{1/2}$ (%)**[e] | 83.5 (33.3) | 99.8 (88.9) |
| **Refinement Statistics** | | |
| **Resolution (Å)** | 38.57-1.60 (1.65-1.60) | 32.0-2.07 (2.14-1.90) |
| **$R_{work}/R_{free}$ (Å)** | 24.4/28.4 | 21.0/23.5 |

| | | |
|---|---|---|
| **Number of reflections in refinement (work/free)** | 31,104/2517 | 7188/1350 |
| **Number of non-H protein atoms** | 1835 | 914 |
| **Number of water molecules** | 218 | 13 |
| **Number of protein residues** | 231 | 116 |
| **RMS (bonds)** | 0.006 | 0.003 |
| **RMS (angles)** | 0.829 | 0.55 |
| **Ramachandran: favored, outliers (%)** | 98.2, 0 | 96.5, 0.9 |
| **Clashscore[f]** | 4.68 | 1 |
| **Wilson B (Å$^2$)** | 21 | 40 |
| **Average B (Å$^2$)** | 28 | 57 |
| **Protein** | 27 | 57 |
| **Chain A** | 24 | 59 |
| **Chain B** | 30 | N/A |
| **Water** | 36 | 53 |

**Table S1.** X-ray data collection and refinement statistics for A120S and A120L.

[a] Numbers in parentheses are for highest resolution shell.
[b] $R_{merge} = \Sigma_{hkl} \Sigma_{i=1,n} |I_i(hkl) - \langle I(hkl) \rangle| / \Sigma_{hkl} \Sigma_{i=1,n} I_i(hkl)$
[c] $R_{meas} = \Sigma_{hkl} \surd(n/n\text{-}1)\ \Sigma_{i=1,n} |I_i(hkl) - \langle I(hkl) \rangle| / \Sigma hkl\ \Sigma_{i=1,n} I_i(hkl)$
[d] $R_{pim} = \Sigma_{hkl} \surd(1/n\text{-}1)\ \Sigma_{i=1,n} |I_i(hkl) - \langle I(hkl) \rangle| / \Sigma hkl\ \Sigma_{i=1,n} I_i(hkl)$
[e] $CC_{1/2}$ = Pearson Correlation Coefficient between two random half datasets.
[f] Number of unfavorable all-atom steric overlaps ≥ 0.4. per 1000 atoms.

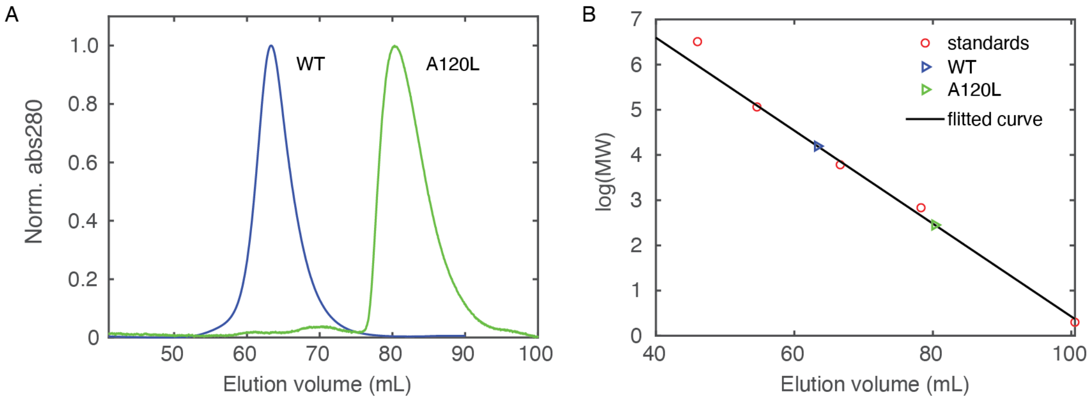


**Figure S1.** (A) The size exclusion chromatography (SEC) Superdex 75 elution profile of WT human TTR and the A120L variant. (B) The calibration curves of the Superdex 75 column. The SEC-measured molecular weights of the WT tetramer and A120L variant are 66.4 and 11.6 kDa, respectively. For comparison, the calculated molecular weight of the WT tetramer and the A120L monomer is 55.6 and 13.9 kDa, respectively. The in-column concentration of both constructs was around 1 μM.

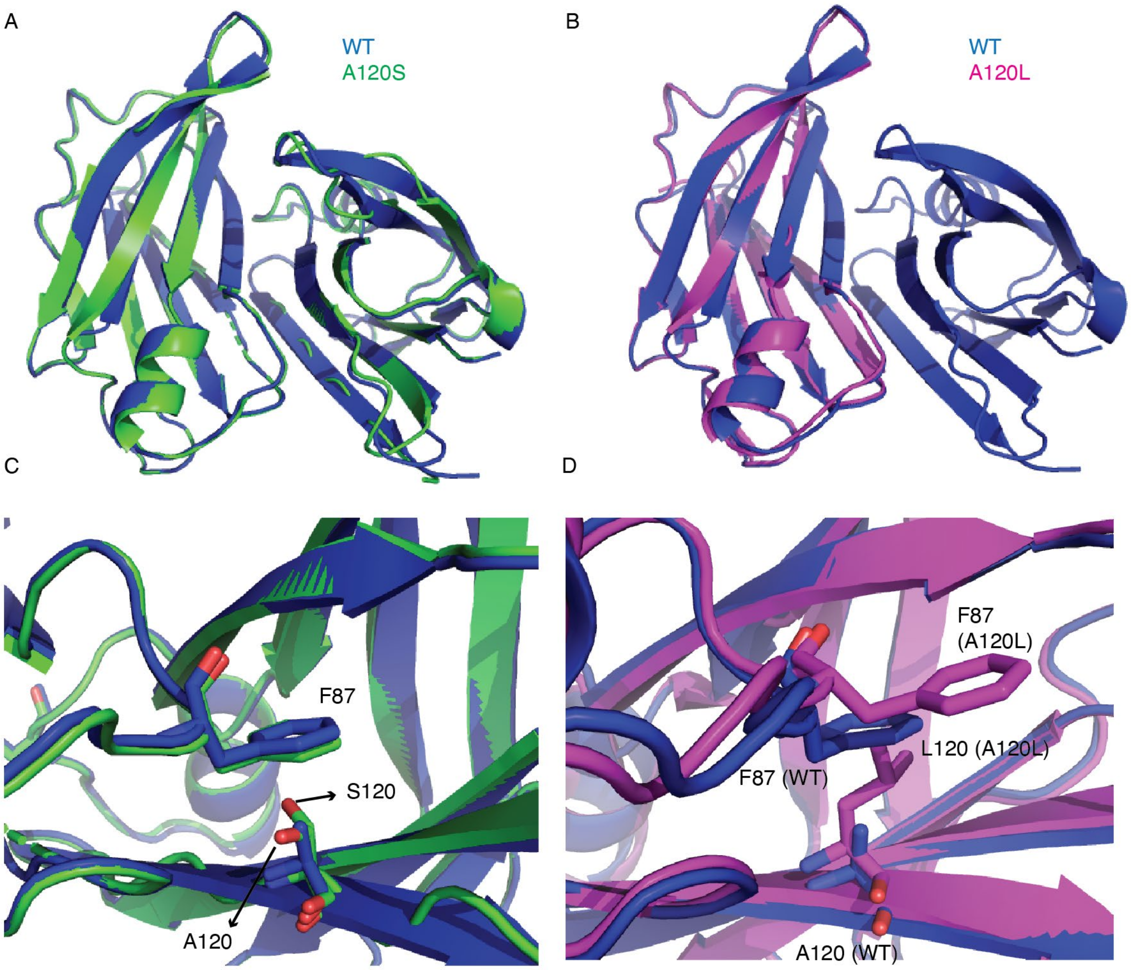


**Figure S2.** The overlaid X-ray structures for WT (blue, PDB: 5CN3) vs. A120S (green, PDB: 8W1N) (A) and WT vs. A120L (magenta, PDB: 8W2W) (B). (C) Close-up views highlighting the F87-A120 interaction for WT TTR (blue) and. A120S (green). Note that there are two distinct orientations of the S120 sidechain. (D) Position of the F87 side chain in WT TTR (blue) and A120L TTR (magenta), where it is displaced from its binding pocket in the neighboring subunit by steric clash with the bulky L120 sidechain. The subunit shown in blue is WT TTR (A120 sidechain). The A120L subunit with L120 shown in sticks is aligned to the WT TTR subunit with A120 shown in sticks.

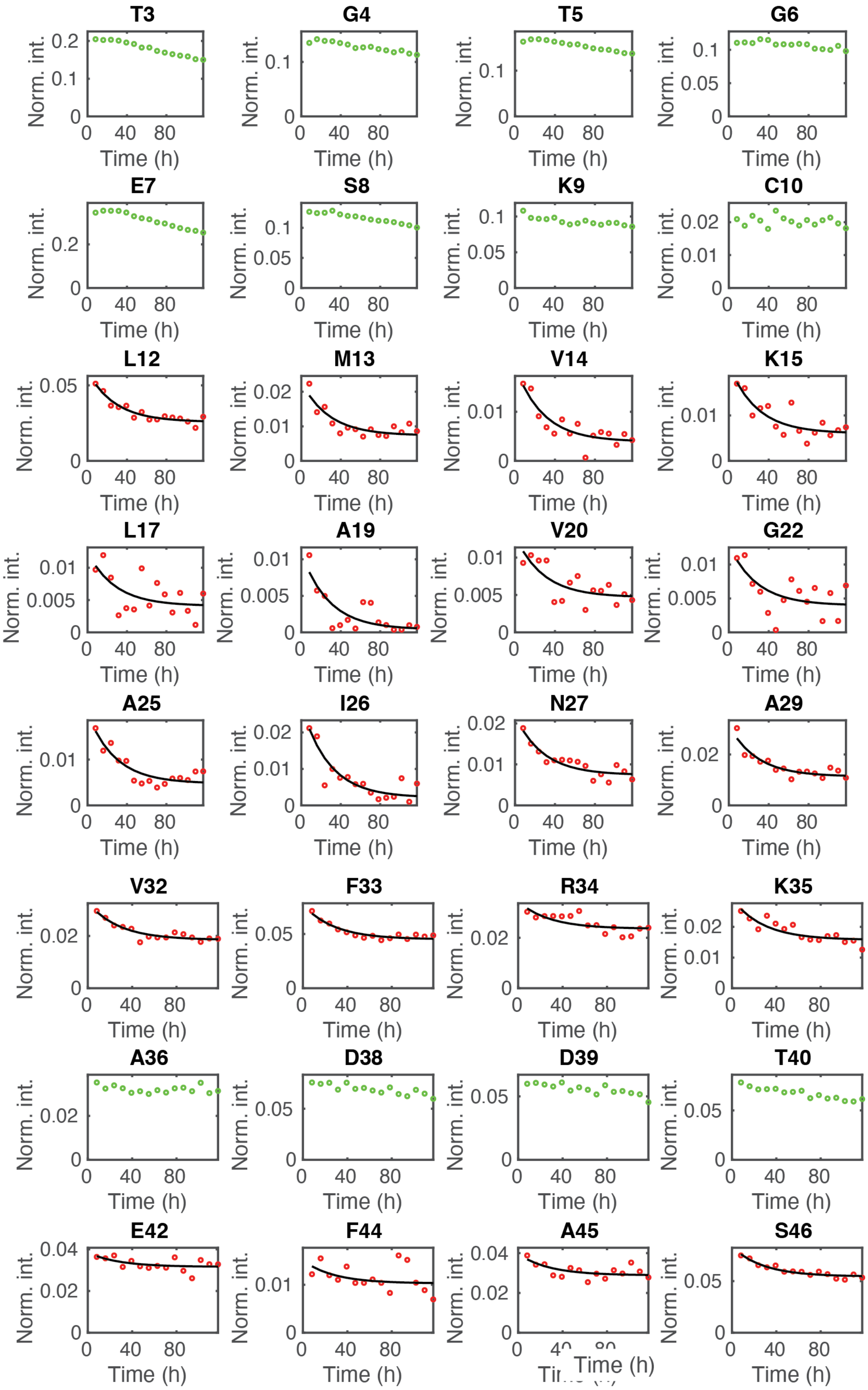
T3
G4
T5
G6
E7
S8
K9
C10
L12
M13
V14
K15
L17
A19
V20
G22
A25
I26
N27
A29
V32
F33
R34
K35
A36
D38
D39
T40
E42
F44
A45
S46
Norm. int.
Time (h)

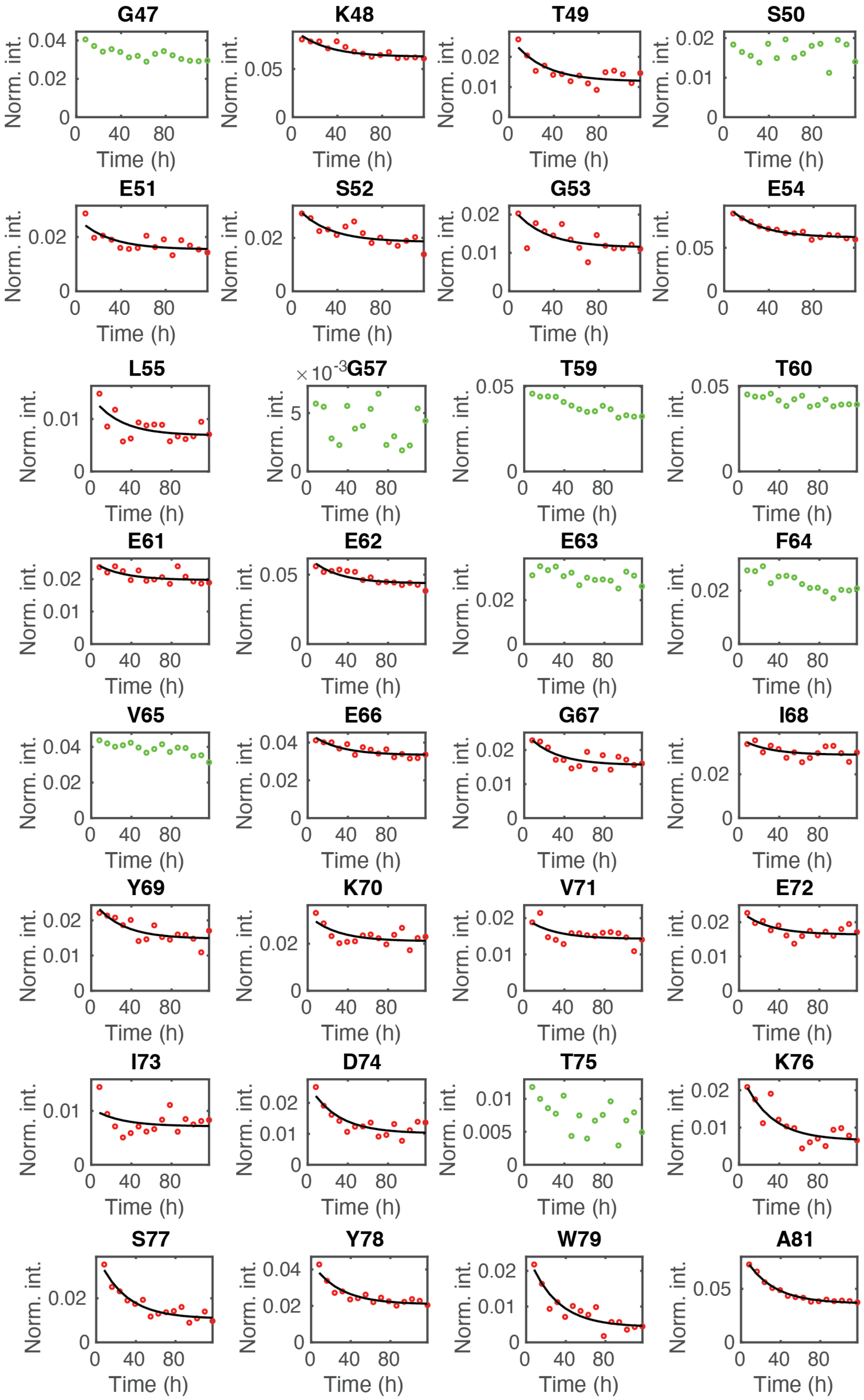
G47
K48
T49
S50
E51
S52
G53
E54
L55
G57
T59
T60
E61
E62
E63
F64
V65
E66
G67
I68
Y69
K70
V71
E72
I73
D74
T75
K76
S77
Y78
W79
A81
Norm. int.
Time (h)

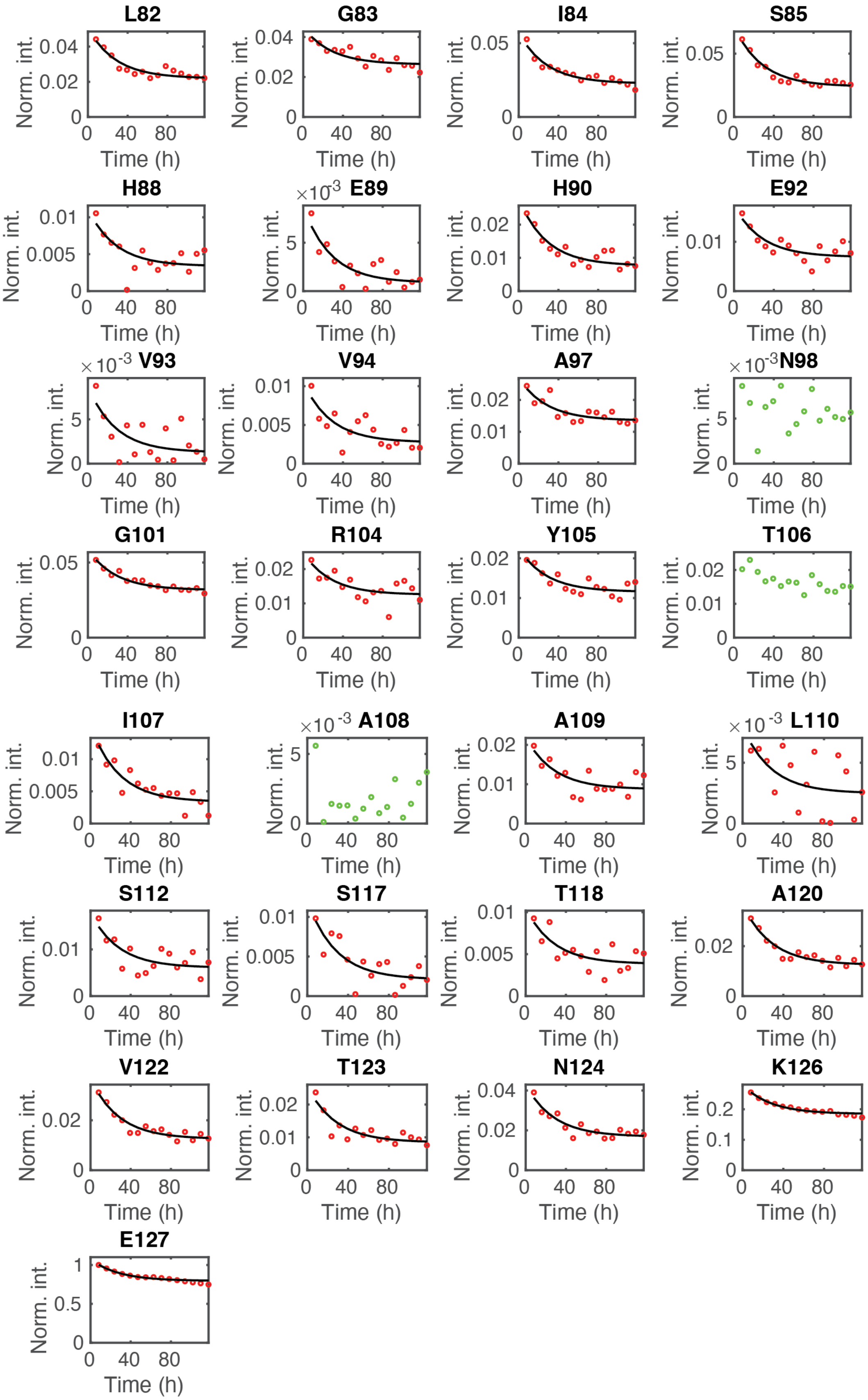
L82
G83
I84
S85
H88
×10^-3 E89
H90
E92
×10^-3 V93
V94
A97
×10^-3 N98
G101
R104
Y105
T106
I107
×10^-3 A108
A109
×10^-3 L110
S112
S117
T118
A120
V122
T123
N124
K126
E127
Norm. int.
Time (h)
0
40
80

**Figure S3.** Time-dependent loss of $^1$H, $^{15}$N cross-peak intensity in spectra of a 1:1 mixture of A120L:$^{15}$N-WT TTR at 298 K and pH 7.0. Residues whose cross-peak intensities in HSQC spectra remain nearly constant are labeled in green and were not included in the single exponential fitting. Data points for residues whose cross-peak intensities show time-dependent decay are shown in red and fitted by a global single exponential function in black solid lines. The peak intensity of the C-terminus E127 was used for normalization.
